# Is the hydrogen of the 2’-hydroxyl group of HENMT1 a *Schrödinger’s* hydrogen?

**DOI:** 10.1101/2023.05.22.541725

**Authors:** Philipp Kaldis, Li Na Zhao

## Abstract

piRNAs are important in protecting germline integrity. 3’-terminal 2’-O-methylation is essential for piRNA maturation and to protect it from degradation. HENMT1 carries out the 2’-O-methylation, which is of key importance for piRNA stability and functionality. However, neither the structure nor the catalytic mechanism of HENMT1 have been studied. We have constructed a catalytic-competent HENMT1 complex using computational approaches, in which Mg^2+^ is primarily coordinated by four evolutionary conserved residues, and is further auxiliary coordinated by the 3’-O and 2’-O on the 3’-terminal nucleotide of the piRNA. Our study suggests that metal has limited effects on substrate and cofactor binding but is essential for catalysis. The reaction consists of deprotonation of the 2’-OH to 2’-O and methyl transfer from SAM to the 2’-O. The methyl transfer is spontaneous and fast. Our in-depth analysis suggests that the 2’-OH may be deprotonated before entering the active site or it may be partially deprotonated at the active site by His800 and Asp859, which are in a special alignment that facilitates the proton transfer out of the active site. Furthermore, we have developed a detailed potential reaction scenario and our study indicates that HEN1 is Mg^2+^ utilizing but is not a Mg^2+^ dependent enzyme.

## INTRODUCTION

RNA modification proteins (RMPs) are (i) enzymes that covalently modify RNA molecules (“writers”); (ii) enzymes that reverse these modifications (“erasers”); and (iii) proteins that recognize and selectively bind these modified RNAs (“readers”) [1,2]. RMPs play a myriad of roles in the structural integrity and translational fidelity of RNAs. The mechanisms of adding, removing, and recognizing a chemical group in RNAs has been referred to as epitranscriptomics. Emerging data has suggested that epitranscriptomics is an important indicator in cancer and other diseases, and RMPs have emerged as a new class of therapeutic targets with a burst of research interest in recent years [1,3].

Over 100 types of reversible and dynamic chemical modifications are carried out by RMPs on cellular RNAs [4,5]. In cancer, 27% of all known human RMPs are dysregulated, among them HENMT1 and LAGE3 have been reported to be the two most frequently overexpressed genes across a wide variety of cancer types. They are consistently overexpressed in tumors at various stages of progression, particularly at stages III and IV, and have been suggested to be promising drug targets for anti-tumor therapies [6].

Micro RNAs (miRNAs) are short non-coding RNA molecules, about 21-24 nucleotides (nt) in length, that regulate gene translation through base-pairing complementary to silence or degrade target mRNAs. They are associated with almost every aspect of biological processes, including cell growth, metabolism, inflammation, apoptosis, and the pathophysiology of many diseases [7]. Recent studies have suggested circulating miRNAs play oncogenic roles and can serve as promising diagnostic and prognostic biomarkers for many diseases [8–16].

Hsa-miR-21-5p, as the first-identified miRNA molecule, is a representative, circulating, and typical onco-miRNA [8,13]. It has been extensively investigated in various malignancies, notably in brain cancer, lung cancer, colorectal cancer, pancreatic cancer, breast cancer, gastric cancer, esophageal cancer, and hepatocellular carcinoma [8,9,11,13,17]. Hsa-miR-21-5p post-transcriptionally regulates the expression of multiple cancer-related target genes, such as the phosphatase and tensin homolog (PTEN), the programmed cell death protein 4 (PDCD4), the reversion-inducing cysteine-rich protein with Kazal motifs (RECK), and the signal transducer activator of transcription 3 (STAT3) [18]. Hsa-miR-21-5p overexpression promotes tumor growth, metastasis and invasion, reduces sensitivity to chemotherapy, and is associated with poor survival in various cancers [17,19,20].

The Hua ENhancer (HEN) methyltransferase 1 (HENMT1; C1orf59; EC 2.1.1.n8; UniPort ID: Q5T8I9) has a pronounced role in 2’ O-methylation (2’-Ome) of mammalian P-element-induced wimpy testis-interacting RNAs (piRNAs) which are critical in the early phases of spermatogenesis, and adult male germ cell transposon repression. HENMT1 loss-of-function induces piRNA instability and ultimately leads to male sterility [21,22]. Recently, HENMT1 was shown to be responsible for the methylation of the 3’-terminal 2’-Ome of mammalian miR-21-5p, which is predominant in human non-small cell lung cancer (NSCLC) tissue [23].

piRNAs are well-defined in the male and female germline, with hundreds of thousands of unique piRNAs in mammals (27,700 sequence in piRNAdb.hsa.v1_7_6.fa from piRNAdb.org [24]; 282,235 clustered piRNAs from piRNA cluster database [25–27]; and 667,944 from piRNABank [28]) [29]. The length of piRNAs varies from 21-31 nt among species with a common and predominant 5’ uridine (U) and conserved A at position 10, but with a less defined secondary structure [30]. In somatic cells, piRNA dysregulation has been associated with tumor development and metastasis and has the potential to predict cancer prognosis.

The 2’-O-methylation on the 3’-terminal of a subset of small RNAs is a crucial step for their functional maturation and is prevalent among fungi, plants, and animals, which is achieved by a conserved SAM-dependent RNA methyltransferase, HEN1 and its homologues. Structural studies of plant HEN1 with a 22 nt RNA duplex and the cofactor product SAH revealed that Mg^2+^ is coordinated by both 2’ and 3’ hydroxyls on the 3’-terminal of the 22 nt RNA and four residues (Glu796, Glu799, His800, and His860; corresponding to E133, E136, H137 and H182 in mouse; and E132, E135, H136, and H181 in human) at the active site of the methyltransferase domain [31]. The SAM-binding pocket is formed by five consecutive residues _719_DFGCG_723_ residing adjacently to the catalytic domain of *Arabidopsis* HEN1. Furthermore, the mechanism of 2’-O-methylation has been suggested to be Mg^2+^-dependent for plant HEN1 [31]. The substrate specificity of plant HEN1 is well-defined since it can methylate both microRNA and small interfering RNAs duplexes (miRNA/miRNA* or siRNA/siRNA*), with a preferred length of 21-24 nt, RNA duplexes with 2 nt overhang, and free 2’- and 3’-hydroxyls on the 3’ terminal nucleotide [32,33]. The MTase domain of Drosophila HEN1 is located at its *N*-terminus and biochemical assays indicate that it can methylate small single-stranded RNAs but not double-stranded RNAs [34,35]. Four crystal structures (PDB IDs: 3JWG, 3JWI, 3JWH, and 3JWJ) of the MTase domain of a bacterial homolog of HEN1 demonstrate a unique motif and domain that are specific for RNA recognition and catalysis [36], with the F*X*PP motif being important for substrate binding [37]. The mouse homolog of HEN1 (mHEN1) is expressed predominantly in testis and methylates the 3′ end of piRNAs in vitro [38]. The methylation efficiency for piRNAs with different 3’ end nucleotide are: A (259%) > C (137%) > U (100%) > G (44%) [38]. mHEN1 does not recognize the 5’ end of the substrate and is not particularly specific about the length of RNA substrate [38].

The architecture of human HENMT1 (393 aa) consists of a confirmed MTase domain, which alone cannot confer catalytic activity [37]. However, including the _27_FKPP_30_ motif at the very *N*-terminus, namely the 26-263 region (below we refer to it as MTase region), confers full activity of the Mtase activity in vitro [37]. Unlike plant HEN1, which methylates double-stranded RNA, the mammalian HEN1 methylates only single-stranded RNA [37]. The *C*-terminal domain (CTD, ∼263-393) of HENMT1 lacks homology in the primary sequence, which indicates that the CTD varies substantially across species. However, there is a possibility that the *C*-terminal domain may together with the very *N*-terminus, cooperatively recognize and bind the substrate; or HENMT1 interacts with other proteins to facilitate its localization as was observed for zebrafish HEN1 [39].

The MTase region of human HENMT1 crystallized with SAH (PDB ID: 4XCX) and SAM (PDB ID: 5WY0) is missing the important cofactor Mg^2+^. However, human HENMT1 has been reported to prefer Mn^2+^ over Mg^2+^, similar to bacteria HEN1 [37]. Glu132, Glu135, His136, and His181, corresponding to Glu796, Glu799, His800 and His860 in plant HEN1, were proposed to be responsible for Mg^2+^ binding by (a) manual assertion inferred from sequence alignment in UniPort entry Q5T8I9; and (b) comparison with HEN1 from plant (PDB ID: 3HTX) [40].

Despite the structural and biochemical advance in studying HENMT1, there are still many questions unanswered [37]. This includes, (a) what is the complete list of substrates of HENMT1? Are piRNAs and miR-21-5p the only substrates? and (b) what is the molecular basis of HENMT1 catalytic activity? Here, we aim to use computational biochemical approaches to investigate the mechanisms driving HENMT1 activity.

From our previous work, we have observed that (a) the energy required for the methyl transfer in SAM-dependent methyltransferases is around 8 k_cal_/mol in protein and 13.8 k_cal_/mol in water, which is close to the spontaneous transfer under thermodynamic conditions; and (b) the proton transfer is a rate-limiting step [41]. Transferring this knowledge to plant HEN1 and mammalian HENMT1, we will consider and examine five potential reaction mechanisms for the rate-limiting proton transfer step: (I) 2’-OH is deprotonated before it reaches the active site in the reaction-ready state; (II) a hydroxide present at the active site acts as base to withdraw the proton; (III) His800 acts as a base; (IV) Glu796 acts as a base; and (V) His800 and Asp859, in a special alignment, facilitate the proton transfer out of the active site.

By compiling available structural information of HENMT1, computational modeling, calculating free energies, as well as a detailed analysis and carefully evaluation of possible mechanisms, we have reached the conclusion that the hydrogen from the 2’-OH has the fate as the “*Schrödinger’s cat*”; i.e. it is likely deprotonated before entering the active site; or it is partially deprotonated once entering the active site due to its interaction with the residues (Glu796 and His800) at the active site; or the spatial alignment of Asp796 and His800 may facilitate the transport of the hydrogen out of the pocket. In addition, we propose an equilibrium ordered kinetic mechanism in which SAM and Mg^2+^ bind first prior to the substrate binding.

## MATERIAL and METHODS

### Structure preparation

Since the truncated *C*-terminal domain (residues 666-942) of plant HEN1 and MTase region of HENMT1 are sufficient for the methyltransferase activity, here we used the truncated plant HEN1 methyltransferase domain (HEN1-M) and MTase region of HENMT1. We have built 3 systems: (SI) plant HEN1-M with 2’-OH binding to Mg^2+^ directly; (SII) plant HEN1-M with 2’-OH binding to Mg^2+^ through hydroxide mediated interactions, termed as HEN1-W; (SIII) mammalian HENMT1 with 2’-OH binding to Mg^2+^ through hydroxide mediated interactions. 3HTX.pdb was the primary template for HEN1-W and HEN1-M. SAH was converted back to SAM by adding a methyl group using Avogadro (Figure S1). The missing loops (_839_TPETQEENNSEP_850_ and _912_SVENV_916_) were built and refined using LoopModel and DOPE LoopModel modules within MODELLER10 [42] while the top 10 out of 400 models were optimized and relaxed within MOLARIX-XG.

For human HENMT1, the canonical sequence 26-263, featuring the MTase domain, was taken from uniprot (ID Q5T8I9). The crystal structure of the human MTase region of HENMT1 in complex with SAH (PDB ID: 4XCX; covering sequence 26-170, 180-232, and 244-262) and SAM (PDB ID: 5WY0; covering sequence 31-85, 105-173, 177-236, and 245-258) were used as templates for modeling the MTase region of the HENMT1 complex with cofactor SAM [see Figure 1(A)]. The automodel and loopmodel modules within MODELLER 10 were used to build 200 models and loop refinement. The final models were selected based on the objective function; the four residues (Glu132, Glu135, His136, and His181) that directly bind Mg^2+^ are refined based on plant HEN1 (3HTX.pdb). Note that the crystal water (W1) from 5WY0.pdb was kept for SII and SIII. The structures were further optimized and relaxed by MD simulations within the MOLARIX-XG package before the free energy calculation.

**Figure 1:**
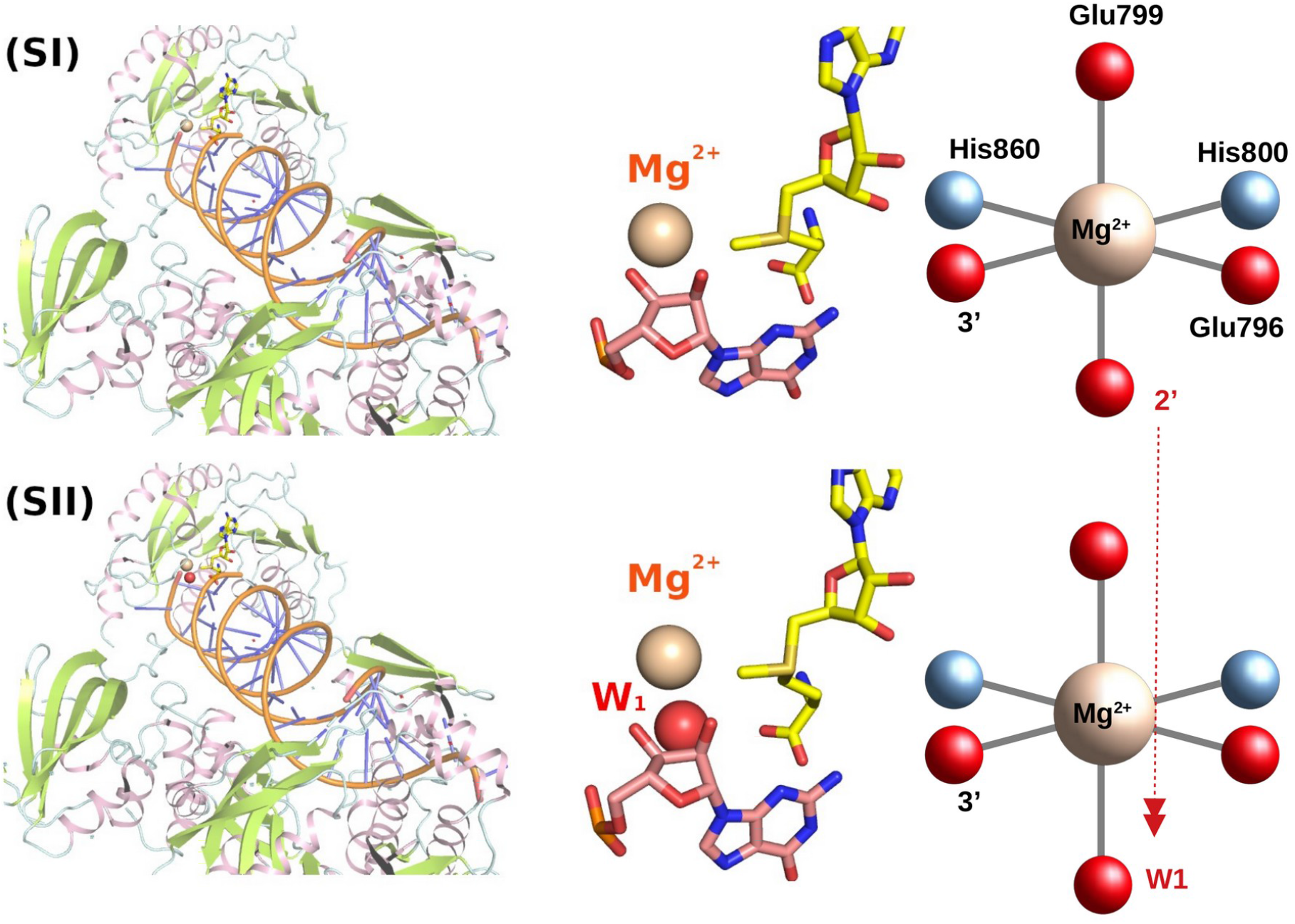
Plant HEN1 with 2’-O and 3’-O groups directly coordinating Mg^2+^ (SI). Plant HEN1 with hydroxide and 3’-O groups coordinating Mg^2+^ (SII).

For plant HEN1, the miR173/miR173* duplex, crystallized in 3HTX, was experimentally generated using synthesized miR173 and miR173* RNA oligonucleotides with 5′ P and 3′ OH annealed [32]. Note that the 2nt 3′ overhang is an important feature of the substrate. The configuration for the miR173/miR173*(22A), miR173/miR173*(22C), miR173/miR173*(22U) are generated by directly mutating the 22^ed^ nucleotide. The detailed sequences are given in Table S3. For plant HEN1, the configurations with the lowest binding energy towards the substrate were selected and used for the subsequent reaction calculation.

### Binding energy calculations

The binding energy calculations were carried out to examine (a) the truncated HEN1 (HEN1-M), and (b) the role of the Mg^2+^. We calculated the binding energy of SAM and substrate, which is a small RNA duplex, derived from one natural substrates of plant HEN1, termed miR173/miR173* [31], in the presence/absence of Mg^2+^ using the Linear Response Approximation (LRA) version of Protein Dipoles Langevin Dipoles (PDLD/S-LRA) method and also its PDLD/S-LRA/β Version [43] within MOLARIS-XG package. At first, we generated both complexes with full length HEN1 (HEN1-FL) and truncated HEN1 featuring the methyltransferase domain (HEN1-M) with and without Mg^2+^ configurations, and with the charged and uncharged forms of solute, respectively, and then treated the long-range interaction with the local reaction field (LRF) [44]. After explicit all-atom molecular dynamics simulations of all above complexes, each lasting 2ps, with the surface-constrained all-atom solvent (SCAAS) [45], then we carried out the PDLD/S calculations on the generated configurations. We took the average value as the consistent estimation of the binding free energy. A 2ps run was done for each of these simulations at 300K. The philosophy behind this method has been discussed in Ref. [43] and in our previous work [41].

### Simulations

The 2’-O-methylation requires two steps: deprotonation of the 2’-OH group precedes the transfer of the methyl group from SAM to 2’-O group. First, we started from the methyl transfer from SAM to 2’-O and later addressed the deprotonation of the 2’-OH group. The initial kinetic descriptions were done at the M06-2X/6-31++G(d,p) level which provide a fine balance between the computational costs and the reliability of results with a continuum solvent model. Mg^2+^ and the surrounding ligands (2’-O/H_2_O/OH^-^, 3’-O, Glu796, Glu799, His800 and His860) were included in the QM region (consisting of 83 atoms). The results from the DFT calculation are then used to calibrate empirical valence bond (EVB) parameters for the methyl transfer in enzymes, both plant HEN1 and HENMT1. The active site region was immersed in a 32Å sphere of water molecules using the surface-constraint all-atom solvent (SCAAS) type boundary condition [45]. The geometric center of the EVB reacting atoms was set as the center of the simulation sphere. The Langevin dipoles was applied outside of this 32Å region, followed by a bulk continuum. The long-range electrostatics were treated with the local reaction field (LRF) method. Atoms beyond the sphere were fixed at their initial positions and no electrostatic interaction from outside of the sphere was considered. In order to determine the protonation state and optimize the charge distribution of all ionizable residues, we were using the Monte Carlo proton transfer (MCPT) algorithm which simulates the proton transfer between charged residues, the charge distribution was updated and evaluated with Monte Carlo approaches to identify the optimal charge distribution. The protonation state of the ionizable residues are shown in Figure S2 in the Supporting Material. The detailed EVB simulation procedures are described in our previous work [41,46,47]. The EVB simulation of the methyl transfer were done using the Enzymix module within the MOLARIS-XG package [45,48]. Note that all the DFT calculations were done using Gaussian 16 Revision C.01 [49].

## RESULTS

### Overview of modeled structure

The overview of modeled HENMT1 is shown in Figure 2A. After detailed structural analysis, we believe that Glu132, Glu135, His136, and His181 constitute the predominant motif involved in binding Mg^2+^ which can then attract 3’-terminal miRNAs or piRNAs by forming a hydrogen bond with the 3’-OH group of the ribose moiety (Figure 2B). In addition, the 2’- and 3’-hydroxyl groups of piRNA bind Mg^2+^ directly in the presence of SAH (Figure 2B).

**Figure 2:**
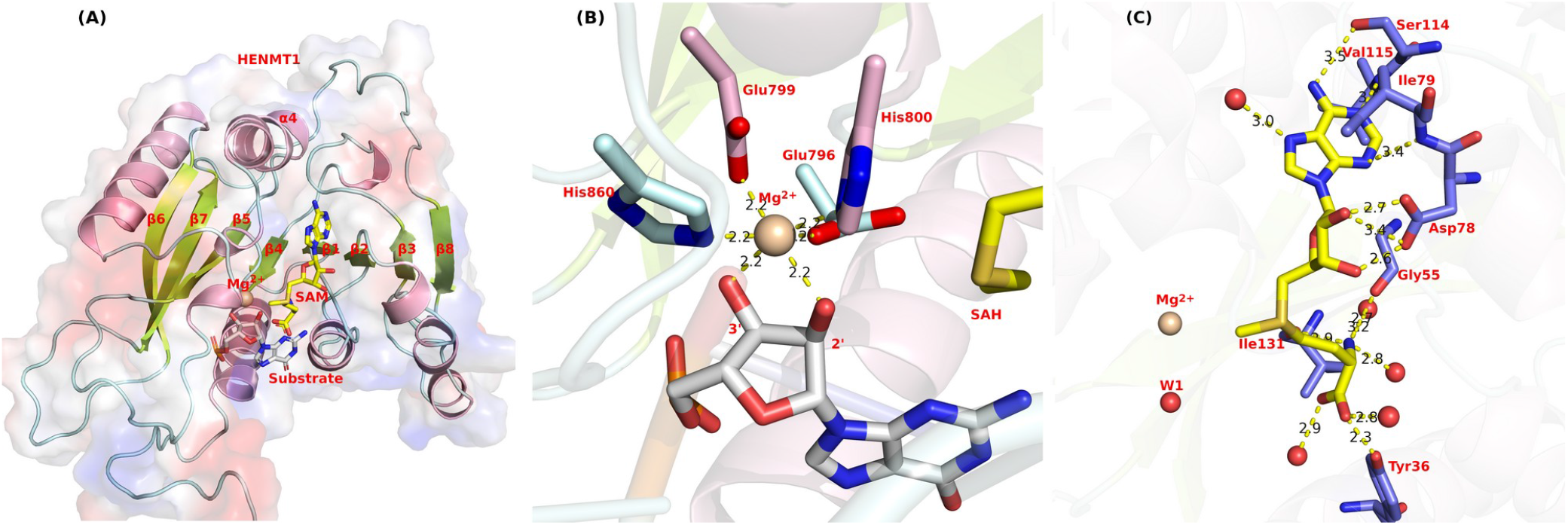
(A) Overview of modeled HENMT1 structure. (B) The Mg^2+^ binding pocket from plant HEN1 (PDB ID: 3HTX). Helices are colored as pink for the C atoms, palecyan for the residues from the loop, and lemon color for the C atoms from β-sheet. (C) The SAM binding pocket (PDB ID: 5WY0) with Mg^2+^ superimposed from 3HTX. The water molecule (W1) is crystal water from 5WY0.pdb.

Plant HEN1 protein not merely has a putative dsRNA binding motif at the *N*-terminus, but also has a conserved SAM-binding motif in the *C*-terminal region [2,32]. Here, we have systematically examined this SAM binding pocket. There are two peptide motifs conserved among plant, mouse, and human isoforms of the canonical SAM binding site; namely, _717_LVDFGCG_723_ and _743_GVDI_746_ in plant HEN1; _52_VADLGCG_58_ and _77_GVDI_80_ in mouse HEN1; and _51_VADLGCG_57_ and _76_GVDI_79_ in human HENMT1.

Figure 2C depicts the interaction scenario of SAM with the HENMT1 in detail. The adenine base of SAM is coordinated by the side chain of Ser114, the backbone of Val115 and Ile79, and a water molecule. The ribose group of SAM is coordinated by equivalent bidentate hydrogen bond interactions between its hydroxyl and the carboxyl group of Asp78. The amino group of methionine is forming a hydrogen bond with the backbone of Gly55 and Ile131, and two water molecules; while the carboxyl group of methionine is forming a hydrogen bond with side chain of Tyr36 and two water molecules. Taken together, we have identified residues 55, 78, 79 from conserved motifs that are directly binding SAM, and are conserved among plant HEN1, animal HEN1, and HENMT1.

### 2’-OH deprotonation

The 2-O methylation requires two steps: (a) the deprotonation of the 2-OH group, and (b) the transfer of the methyl group from SAM to the deprotonated 2-O group. The deprotonation is the prerequisite and it is usually elusive. Since this is hard to determine with theoretical or empirical methodological knowledge, we are trying various different mechanisms to determine which one is the most likely.

We have considered five potential mechanisms: Mechanism I is based on deprotonation during the process when substrate was recruited at the active site. Our calculations, however, determined that the 2’-OH of the ribose sugar has a high instinct p*K*_a_ (12∼14), which is usually neutral at biological pH [50]. Hence, we have considered the possibility that 2’-O(H) still has the H once it enters the reactive site, which needs a base to remove this H. Mechanism II is based on a hydroxide/water present at the active site of HENMT1 crystal structure (5WY0) to deprotonate the 2’-OH. Mechanism III is based on His800 deprotonating the 2’-OH group. Nevertheless, scrutinizing the crystal structure of HEN1 (3HTX): we found that the pKa of His800 (corresponds to His136 in HENMT1) is calculated to be 10.2, which indicated that it binds proton tightly, increasing the likeliness that the hydrogens are on both *δ* and ε sites at product state, supporting a mechanism, in which His800 deprotonates the 2’-OH. Mechanism IV is based on Glu796 deprotonating the 2’-OH group. However, the unusually low pKa value of Glu796 which is estimated to be 1.02 and Glu796 is in the immediate vicinity of 2’-OH. Mechanism V is based on structural observations concerning the spatial alignment of His800 and Asp859, as well as surrounding residues facilitating a proton transfer network to deliver the proton out of the active site. Specifically, Asp859 abstracts a hydrogen from His800 at *δ* site and His800 withdraws the hydrogen from the 2’-OH group to its ε site.

#### Schrödinger’s hydrogen

There are several possibilities for the mechanism that the hydrogen on the 2’-OH group may be lost or still bonds in the process of substrate recruitment to the active site. One possibility is that the 2’-OH group was deprotonated during its recruitment into the active site. Once at the active site, 2’-OH may be (a) already deprotonated; (b) partial deprotonated; or (c) still protonated. For the possibility that 2’-OH needs to be deprotonated at the active site, we started to examine the potential orientation of this hydrogen because this determines the potential base that will extract this hydrogen. One approach is to infer from the potential occurrence of hydrogen bonds based on the positions of proximity heavy atoms from the X-ray structures. This inference of precise orientation of the 2’-O-H bond is of great importance in clarifying the elusive step of the 2’-OH deprotonation by revealing to which heavy atom the proton will be transferred to.

There are two possible orientations of hydrogen of the 2’-hydroxyl group based on the hydrogen bond geometry criteria. The possible hydrogen bonds are illustrated in Figure 3(a), and the possible orientation of the hydrogen either towards (i) O4’ of the same sugar [Figure 3(b)], or (ii) away from the O4’ of the same sugar [Figure 3(c)].

**Figure 3:**
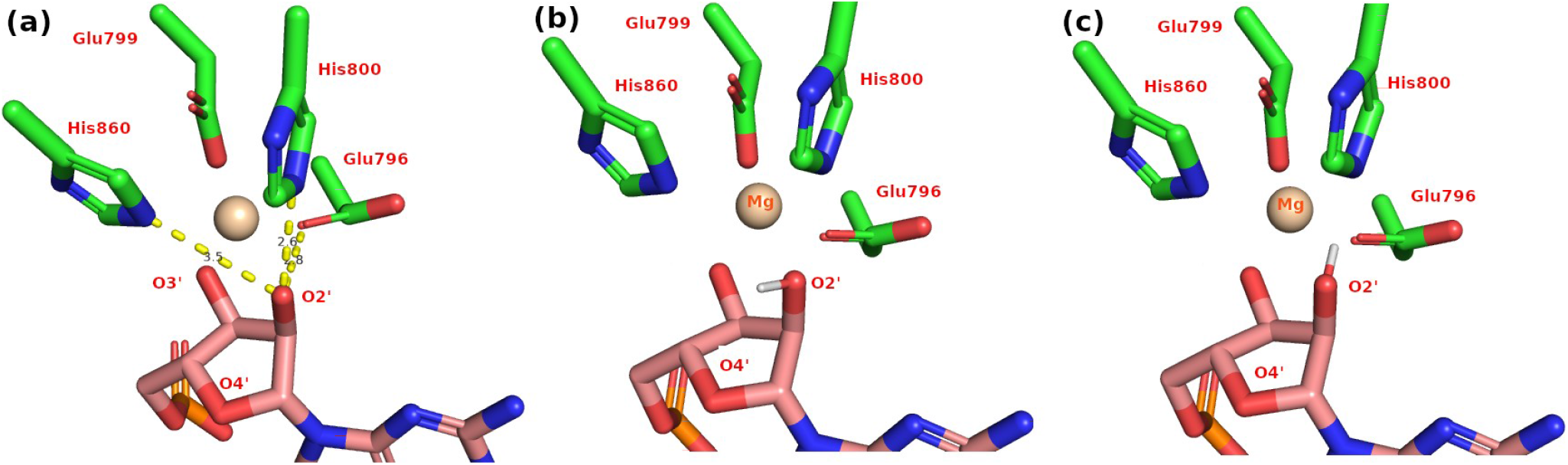
The potential hydrogen bonds and hydrogen orientations. (a) The potential hydrogen bond between the 2’-OH group and the surrounding atoms. (b) Hydrogen orientates towards the O4’ of the same sugar related to the hydrogen bond with His860. (c) Hydrogen away from the O4’ of the same sugar, related to the hydrogen bond forming with His800 and Glu796.

When hydrogen is orientated towards the direction of Glu796 and His800, our QM calculation of the active site indicate that O2’ is partially deprotonated (see Figure S5), while Glu796 and 2’-O share the hydrogen with d_02’-H_ = 1.02Å and d_Glu796-O_ = 1.58Å. Because of this, the orientation of the hydrogen at O2’ is essential to know.

#### Presence of water/hydroxide at the active site

In order to understand the chemical reactions, we have calculated the pKa value of the residues at the active site using both PropKa and MCPT approaches. The pKa value of residues His800, His860, Glu796, and Glu799 were calculated based on the plant HEN1 crystal structure (3HTX.pdb) using PropKa [51] and found to be 10.2, 5.3, 1.0, and 7.7, respectively. Note that the crystal structure 3HTX was obtained at pH 4.8. Based on the pKa calculation, we conclude that His800 is a strong base and strongly bound to the positively charged Mg^2+^. Glu796 and Glu799 are negatively charged; the lower pKa value of Glu796 may also be due to its binding to Mg^2+^. A more thorough calculation using MCPT confirms the pKa value calculated by PropKa. Since 3HTX is obtained with pronounced product SAH, we postulate that His800 and Glu796 are at the product state, which implies a possible mechanism with His800 withdrawing a proton. Even His800 may help deprotonate the 2’-OH for the subsequent methyl transfer, but it won’t be energy favorable to replace the hydrogen and Mg^2+^ that tightly binds the N_δ_ and N_ε_ with the hydrogen from 2’-OH.

Since Mg^2+^ is positively charged, it would be electrostatically favorable to bring the substrate with 2’-O rather 2’-OH in close proximity towards Mg^2+^. If 2’-OH is deprotonated *in situ*, an external base must be coordinated by Mg^2+^. Since Mg^2+^ already has four firm coordination atoms, together with the incoming 2’-O(H) and 3’-OH group from the substrate, resulting in a perfect six coordination as preferred by Mg^2+^, indicating a lack of right coordination for an external base. Therefore, such an external base may be present during the process when the substrate was recruited to the active site. The external base with a high likelihood to be a hydroxide may abstract a proton from 2’-OH, which was made further possible because of a heavy O atom was crystallized near the active site before the substrate was recruited in the HENMT1 complex with SAM (5WY0.pdb). We have excluded that the heavy O atom is water because the energy barrier of deprotonation of water is estimated to be 29 *k*_*cal*_*/mol* [52], which makes this reaction pathway not energy feasible. Meanwhile, as an earlier study has shown that the energy cost to transfer a hydroxide from the bulk solution to the Mg^2+^ coordination shell is about 5 k_cal_/mol [52], and the free energy of hydroxide ion formation based on *k*_*B*_*T* ln(10)(15.7−7.4) is 11.3 *k*_*cal*_*/mol* at pH 7.4. Therefore, the hypothesis that a hydroxide is present is more feasible than a water for the reaction to take place.

The catalytic cycle starts from hydrated metal [termed as M; Mg^2+^(H_2_0)_6_ or Mn^2+^(H_2_0)_6_] binding to HEN1 forming a binary complex (E·M) with the replacement of 4 water molecules. SAM binding to HEN1 has no direct contact with metal. During RNA binding, it is possible to have the 3’-O of the ribose moiety from the substrate to replace one O from the inner sphere of Mg^2+^/Mn^2+^, however, it is not energetically favorable to have the 2’-OH group of the substrate to replace the last O atom at the inner sphere. The reason is that (a) the remaining O provides a base to abstract the proton from 2’-OH for the proceeding methyl transfer since no other base is available in the immediate vicinity; and (b) it is electrostatically favorable for a positively charged methyl group (-CH_3_) to transfer to 2’-O that is directly coordinated by water rather than the positively charged Mg^2+^. If the 2’-OH binds Mg^2+^ directly, it is not electrostatic favorable for the methyl group (-CH_3_^+^) to come to the 2’-O group which has +2 charge in the immediate vicinity. In addition, a base is required to deprotonate the 2’-OH for the forthcoming methyl transfer, since the only two potential base residues (Glu796 and His800 in HEN1; Glu132 and His136 in HENMT1) at the active site are directly bound to the Mg^2+^ (equal to 2 protons) which is not electrostatic favorable to withdraw another proton from 2’-OH group. Recently a water/hydroxide (W1 in see Figure 2C) in the active site of human HENMT1 was reported [37]. Therefore we first assumed that (a) W1 instead of 2’-OH provides the direct binding to Mg^2+^ and (b) W1 as hydroxide withdraws the proton from 2’-OH, which also solves the piece of the puzzle lacking a base to deprotonate the 2’-OH group. However, W1 exists in the active site which is lacking metal and substrate (5WY0.pdb). It is highly likely once the metal and substrate binding happened, the water won’t be there anymore, which is evident in 3HTX.pdb. Meanwhile, we have tried the water flooding approach (see Supporting Material) in an attempt to saturate the active site with water and found it is not possible to insert water. Furthermore, we were unable to identify the transition state for this speculated proton transfer to happen. Therefore, we have excluded the possibility of a hydride/water at the active site acting as a base to facilitate the 2-OH deprotonation. Cumulatively, our above structural analysis provides a foundation for our catalytic mechanism study.

### The role of Mg^2+^

Mg^2+^ possesses a strict octahedral geometry with a coordination number 6. Initial scrutinizing the crystal structure of plant HEN1 (PDB ID: 3HTX), we find that Glu796, Glu799, His800, and His860 together with the 2’- and 3’-OH of the substrate RNA, fulfill the coordination requirement of Mg^2+^, and Mg^2+^ plays a direct structural role at active site. However, this structure does not reflect the reaction scenario since it is a chimera of product and reactant: (a) product SAH at the pocket instead of reactant SAM; and (b) reactant substrate at the pocket but lacks of base to abstract the proton from the 2’-OH for the subsequent 2’-O methylation. Even though it is not a catalytic competent conformation, 3HTX provides a hallmark structure for studying HEN1.

In order to examine the role of magnesium binding, we constructed a system without Mg^2+^, in which the protonation state of residues (Glu796, Glu799, His800, and His860) that previous coordinated the Mg^2+^ are re-evaluated and obtained similar results as the presence of Mg^2+^ (see Table S1). Furthermore, the protonation state of His800 and His860 are both reasonably assigned on the *δ* site to facilitate Mg^2+^ binding.

The role of Mg^2+^ in terms of binding was calculated using the Linear Response Approximation (LRA) version of Protein Dipoles Langevin Dipoles β version (PDLD/S-LRA/β)^35^ within the MOLARIS-XG package. The averaged binding energy is shown in Table 1. Mg^2+^ does not enhance the positively charged cofactor SAM binding, on the contrary, it is not favorable for SAM binding. For the negatively charged substrate RNA, Mg^2+^ did not significantly enhance the substrate RNA binding. From above calculations, we have excluded a positive role of Mg^2+^ in both cofactor and substrate binding.

**Table 1:** The averaged absolute binding energy calculation for SAM and RNA with (w/) and without (w/o) Mg^2+^ for plant HEN1. Note that the unit is k_cal_/mol.

| Systems |  |  | Binding energy |
| --- | --- | --- | --- |
| SAM | w/ RNA | w/ $Mg^{2+}$ | 39.51 |
| | | w/o $Mg^{2+}$ | 25.39 |
| | w/o RNA | w/ $Mg^{2+}$ | 34.19 |
| | | w/o $Mg^{2+}$ | 13.81 |
| RNA | w/ SAM | w/ $\text{Mg}^{2+}$ | -22.31 |
| | | w/o $\text{Mg}^{2+}$ | -23.76 |
| | w/o SAM | w/ $\text{Mg}^{2+}$ | -25.82 |
| | | w/o $\text{Mg}^{2+}$ | -23.11 |

The properties of Mg^2+^ in the inner coordination sphere feature a very tight interaction to either water or proteins. In the case of HEN1, the inner sphere coordination of Mg^2+^ was primarily bound by four residues from protein; and its interaction with the substrate would be with the 2’-OH and 3’-OH of RNA. We studied the electrostatic role of Mg^2+^ during the methyl transfer by calculating the sum of the partial charges of the atoms [shown in Figure 4(A)]. However, the charge of the magnesium does not change much during the methyl transfer stage (Figure 4(B) and Table S2).

**Figure 4:**
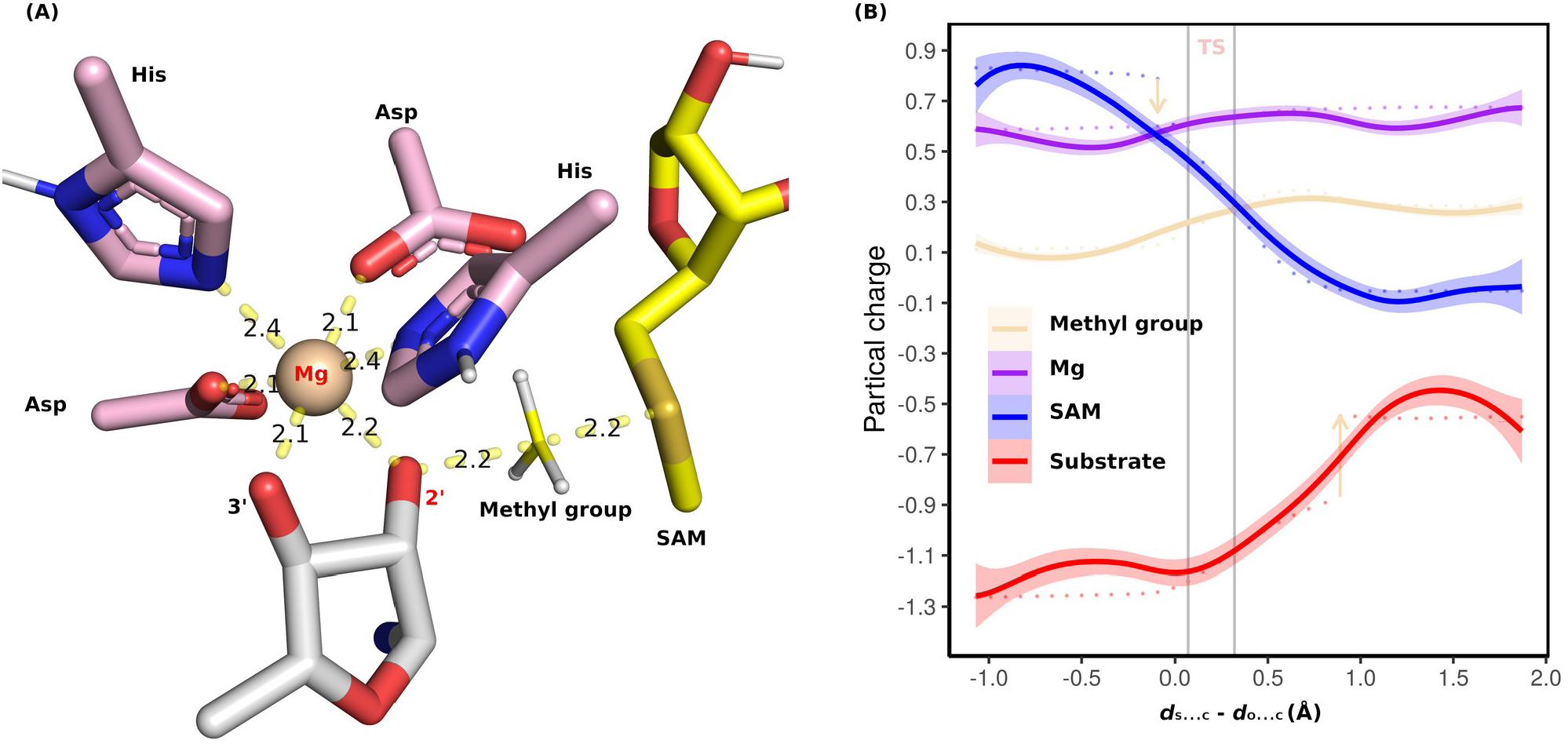
(A) A snapshot of the transition state (TS) for the methyl transfer. All the atoms shown above are included in our QM region. (B) Partial charge across the reaction coordinate. The x-axis indicates the distance of the reactive S from SAM and C from the methyl group minus the distance between the reactive 2’-O and the C of the methyl group. Sum of partial charges on SAM (blue), 2’-O substrate (red), methyl group (lightyellow), and Mg (purple) are indicated by lines. The down-facing arrow indicates that the charge of the methyl group starts to drift away from the SAM group, while the up-facing arrow indicates that the charge of the methyl group starts to be incorporated in the 2’-O substrate. The region between the two gray vertical bars is the TS.

### HEN1 activity is Mg^2+^ utilizing but not Mg^2+^ dependent

First, we examined the current metal (Mg^2+^ or Mn^2+^) binding pocket within the protein. Based on well-established knowledge about plant HEN1, we propose a couple of interaction scenarios for HENMT1 because the metal binding pocket is conserved in both plant HEN1 and human HENMT1. For HENMT1, the MTase region consists of the well-conserved seven-stranded β-sheet [53] and one extra β-sheet, in the order of 67541238, sandwiched between helices [see Figure 2(A)]. Glu132, Glu135, and His136 are located in a short helix between β4 and α4; and His181 is located in another short helix after β5. The two short helices are two unique features in plant HEN1, which are not conserved in RNA and DNA MTases [36].

Second, we will examine the Mg^2+^ cofactor in HEN1 since it has been proposed that HEN1 catalytic activity is Mg^2+^ dependent [31]. Since it is well known that the distance between Mg^2+^ and its coordination oxygen atom from proteins or small molecules is around 2.07Å which was determined by crystal structures [54]. Scrutinizing the distance in the crystal structure of plant HEN1 (3HTX.pdb), we measured the distance between oxygen and metal as 2.2Å. This is outside the ideal range for Mg^2+^. One possible candidate metal is Mn^2+^. As the ionic radius of Mn^2+^ (0.75Å) is slightly bigger than Mg^2+^ (0.65Å) [54,55], and the distance between Mn^2+^ and oxygen is 2.17Å determined by crystal experiments, which is 0.1Å longer than the distance between Mg^2+^ and oxygen [54,56,57]. The measurements of 2.2Å in plant HEN1 crystal structure are in the range of Mn^2+^-O distance [58]. Since Mn^2+^ has the same coordination geometry as Mg^2+^, and the experimental observation of (a) replacement of Mg^2+^ with Mn^2+^ in the Mg^2+^-utilizing enzymes usually does not change the catalytic activity of enzymes while (b) replacing Mn^2+^ with Mg^2+^ in Mn^2+^-dependent enzyme are less often catalytically competent [54]. Hence, we think Mn^2+^ is likely to be the alternative metal in the HEN1 binding pocket. Furthermore, the study of bacterial *Clostridium thermocellum* HEN1 (*Cth*HEN1) indicates the preference for Mn^2+^ over Mg^2+^ [59].

Third, there may be a catalytic advantage for Mn^2+^ instead of Mg^2+^. The optimal coordination geometries of Mg^2+^/Mn^2+^ in proteins is octahedral, with a firm coordination number of 6. The solvent exchange rate for the inner sphere of Mg^2+^ is within 10^−5^ seconds (10^5^ s^-1^), and for Mn^2+^ is around 5×10^6^ s^-1^ [60,61]. Mg^2+^ has a slower solvent exchange rate compared to Mn^2+^ as it has a smaller radius and higher charge density. Glu132, Glu135, His136, and His181 constitute the predominant motif in coordinating Mg^2+^/Mn^2+^ and contribute 4 out of 6 coordination numbers, while the energy penalty to reduce the coordination number from 6 is high [54]. In theory, Mg^2+^/Mn^2+^ would interact with the 2’-OH and 3’-OH through inner sphere coordination, which implies that (a) prior substrate binding, there maybe two water molecules that fulfill the vacancy left by the HEN1; and (b) during substrate binding, a very low rate of ligand exchange and high energy of partial dehydration may happen. As mentioned earlier, the energy penalty for loss of two waters and bound to the substrate is higher for Mg^2+^ than Mn^2+^ [54]. In terms of catalytic activity, Mn^2+^ in the binding pocket may improve the speed of the reaction.

### Kinetics of plant HEN1 methyltransferase activity

Kinetic analysis of the full-length plant HEN1 (HEN1-FL) revealed that the Michaelis constants for microRNA (a synthetic RNA duplex corresponding to miR173/miR173* from *Arabidopsis thaliana*) and cofactor are 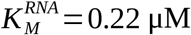 and 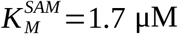, respectively, with an apparent catalytic turnover rate (k_cat_) of 3.0 min^-1^ [62]. The truncated *C*-terminal domain (residues 666-942) of HEN1, termed HEN1-M, is sufficient for methylation with much higher K_M_ values for both RNA 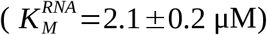 and SAM 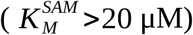, but similar k_cat_ value [62], which indicates that the *N*-terminal residues (1-665) are mainly responsible to enhance the binding of RNA. We have calculated the binding energy of SAM and RNA towards HEN1-FL and HEN1-M, and our results show that HEN1-FL indeed enhances the binding of RNA (see Table 2), which contributes close to 10% of the binding affinity. Further examination of the presence/absence of Mg^2+^ indicates that Mg^2+^ slightly contributes more to enhance substrate RNA binding than the *N*-terminal domain. Hence, our result confirmed that the methyltransfer domain, as a standalone catalytic domain, is the decisive factor for the methyltransferase activity.

**Table 2:** Calculated binding energy (k_cal_/mol) for full-length plant HEN1 and the truncated methyltransferase domain.

| System | $E_{binding}^{SAM}$ | $E_{binding}^{miR173}$ | $E_{binding}^{miR173*}$ |
| --- | --- | --- | --- |
| HEN1-FL (w/ $\text{Mg}^{2+}$) | 39.51 | -28.98 | -27.37 |
| HEN1-FL (w/o $\text{Mg}^{2+}$) | 20.55 | -25.57 | -23.94 |
| HEN1-M (w/ $\text{Mg}^{2+}$) | 40.02 | -26.24 | -26.06 |
| HEN1-M (w/o $\text{Mg}^{2+}$) | 20.26 | -23.98 | -26.97 |

### HENMT1 substrates are not sequence-specific but there is a preference

HENMT1 contains a putative dsRNA binding motif at the *N*-terminus. We have analyzed the direct contact of plant HEN1 and substrate. There are around 46 residues from HEN1 that have direct contacts with less than 3.5Å coordination distance (analyzed from 3HTX.pdb) from substrate. From these, 20 residues establish direct hydrogen bonding contact (see Figure S3). Scrutinizing these contacts, we found that plant HEN1 recognizes and binds substrate RNA primarily through (a) direct binding to oxygen atoms (OP1/2) of phosphate groups; and (b) O2’ and O3’ of the ribose hydroxyl group. There is no direct contact of plant HEN1 with nucleobases, hence, the recognition and binding is not sequence specific, which explains why plant HEN1 has a broad substrate specificity.

The architecture of HENMT1 (393 aa) consists of a confirmed MTase domain, which alone cannot confer catalytic activity,^31^ however, including the _27_FKPP_30_ motif at the very *N*-terminus, namely the 26-262 region (referred as MTase region), confers full activity of the MTase activity.^31^ The current available crystal structures of HENMT1 are 4XCX.pdb and 5WY0.pdb, which cover the majority of sequence from 26-262 (missing 171-179, and 233-243), and 31-258 (missing 86-104, and 237-244), respectively. There are two missing regions; 1-25 at *N*-terminus, and 263-393 at *C*-terminus in the available crystal structures. The incompleteness of the crystal structures limits our investigation regarding the substrate binding mode. Since substrates of animal HEN1 and HENMT1 are predominantly small single-stranded RNAs, which can be explained by the missing dsRNA binding motif in both of them (the sequence alignment can be found in Figure S4). This also provides a logic explanation for the findings [63] in bacterial HEN1: (a) RNA length from 12-24 nt does not affect binding but the activity decreased significantly with 9 nt RNA, and (b) is not sequence specific except the preference for the G at the 3’-end. The knowledge gained from bacterial Hen1 regarding RNA length and RNA sequence specificity may also apply to HENMT1. However, whether the missing regions 1-25 and 31-258 contribute to the substrate recognition or binding is worthy of further study.

## DISCUSSION

The presence of the 2’-hydroxyl (2’-OH) group on the RNA ribose is a distinct feature that differs it from DNA, which facilitates RNA structure folding and enables it to exert profound structural, dynamics and functional characteristics beyond what DNA has. However, the 2’-OH group renders RNA more vulnerable towards degradation and chemical modifications such as methylation and uridylation. For example, the 2′-OH group may nucleophilic attack its adjacent phosphate backbone, resulting in RNA self-cleavage. The 3’-terminal 2’-OH group is subjected to methylation, which in turn protects RNAs from degradation. It has been experimentally challenging to locate the precise position of the hydrogen atom of 2’-OH group or any hydrogen atoms. It is even more challenging to define whether the 2’-O group is protonated or deprotonated at the active site when the local environment involves proteins, metals and is catalytically active. For piRNA, it is crucial to methylate the 2’-O group for its functionality, which is carried out by HENMT1. How the 2’-OH is deprotonated and how the methyl transfer happens is largely unclear. In our work, we have attempted to decode these two processes and the precise position of hydrogen from 2’-OH holds the key to answer the elusive step of the 2’-OH deprotonation. We have carried out extensive discussions and reasoning, and our conclusion is that the hydrogen from 2’-OH is a Schrdinger’s hydrogen, and there are two possibilities for this hydrogen: (a) to be deprotonated before entering the active site or (b) successfully make it to the active site, and may be partially deprotonated by His800 and Asp859, which are in a special alignment that facilitates the proton transfer out of the active site. In summary, there are different possibilities for the deprotonation to happen and the environment acts as hyperparameters to “tune” the reaction.

There are few studies of HENMT1, no structure of HENMT1 with substrates has been published, and the substrates are poorly defined. In our work, we have attempted to model its structure inferred from plant HEN1, and we did a thorough examination of the binding pocket, in which a metal is assumed to be present and catalytically involved. With detailed structural and computational studies, we think HENMT1 can use Mg^2+^ as cofactor, but is not Mg^2+^ dependent. It can also use Mn^2+^ as cofactor and there are advantages to using Mn^2+^ instead of Mg^2+^.

One substantial difference between plant HEN1 and animal HEN1 is the substrate specificity. For plant HEN1, the substrates are double-stranded miRNA or siRNA duplexes, while for animal HEN1, substrates have to be single stranded. In contrast to plant HEN1, that has distinct nucleic-acid-binding channels which implies its substrate has a well defined length (a preference for 21-24 nt RNA with a 2 nt overhangs [32,33]) and distinct loading and cleavage product release. In contrast, animal HEN1, due to lack of pronounced binding grooves, has a broad substrate specificity (piRNAs, Ago2-associated siRNAs, and tRNA-derived sncRNAs) [53] with tolerance for the length of the substrate. For HENMT1, we believe that the substrate is not sequence specific, which is important for its broad substrates (different piRNA). However, HENMT1 may have preferences for substrates like small single-stranded RNAs with certain length, which will need to be studied further in the future.

Although the function of mammalian (mostly mouse) HENMT1 has been firmly established in fertility [37], there are recent reports that HENMT1 may also play an important role in some types of cancer [64]. Based on this and our own results regarding the catalytic activity of HENMT1, we assume HENMT1 plays a role in proliferation in cancer cell lines derived from many different tissues. Our analysis indicates it is associated with breast cancer, lung cancer, and skin cancer (see Figure S6). The role of HENMT1 has been characterized in male fertility, but not yet in females. Similarly, it is possible that HENMT1 is related to the Breast Ductal Carcinoma but this needs further studies.

## Abbreviations

2’O-me: 2’ O-methylation
AGO2: Argonaute RISC Catalytic Component 2
CTD: C-terminal domain
EVB: empirical valence bond
HEN1-FL: full-length plant HEN1
HEN1-M: truncated plant HEN1
HENMT1: Hua ENhancer (HEN) methyltransferase 1
LRA: Linear Response Approximation
MCPT: Monte Carlo proton transfer
miRNAs: microRNAs
NSCLC: non-small cell lung cancer
nt: nucleotides
PDCD4: programmed cell death protein 4
PDLD/S-LRA/β: LRA version of Protein Dipoles Langevin Dipoles β version
piRNAs: P-element-induced wimpy testis-interacting RNAs
PTEN: phosphatase and tensin homolog
RECK: reversion-inducing cysteine-rich protein with Kazal motifs
RMPs: RNA modification proteins
SAH: S-Adenosyl-L-homocysteine
SAM: S-adenosylmethionine
SCAAS: surface-constraint all-atom solvent
sncRNAs: small noncoding RNAs
STAT3: signal transducer activator of transcription 3

## DATA AVAILABILITY

All other data is contained within the manuscript and/or supplementary files.

## SUPPLEMENTARY DATA

Supplementary Data are available at NAR online.

## ACKNOWLEDGEMENTS

The computations were enabled by resources provided by the Swedish National Infrastructure for Computing (SNIC) on resource Kebnekaise at HPC2N partially funded by the Swedish Research Council through grant agreement no. 2018-05973. Additionally, L.N.Z acknowledges the computational support provided by Prof. Warshel’s laboratory at the University of Southern California. L.N.Z is very grateful for the kind support from Prof. Patrik Midlöv.

## REFERENCES

1. Boriack-Sjodin PA, Ribich S, Copeland RA. RNA-modifying proteins as anticancer drug targets. Nature Reviews Drug Discovery. 2018;17: 435–453. doi:10.1038/nrd.2018.71

2. Tkaczuk KL, Obarska A, Bujnicki JM. Molecular phylogenetics and comparative modeling of HEN1, a methyltransferase involved in plant microRNA biogenesis. BMC Evol Biol. 2006;6: 6. doi:10.1186/1471-2148-6-6

3. Shi H, Wei J, He C. Where, When, and How: Context-Dependent Functions of RNA Methylation Writers, Readers, and Erasers. Mol Cell. 2019;74: 640–650. doi:10.1016/j.molcel.2019.04.025

4. Roundtree IA, Evans ME, Pan T, He C. Dynamic RNA Modifications in Gene Expression Regulation. Cell. 2017;169: 1187–1200. doi:10.1016/j.cell.2017.05.045

5. Jonkhout N, Tran J, Smith MA, Schonrock N, Mattick JS, Novoa EM. The RNA modification landscape in human disease. RNA. 2017;23: 1754–1769. doi:10.1261/rna.063503.117

6. Begik O, Lucas MC, Liu H, Ramirez JM, Mattick JS, Novoa EM. Integrative analyses of the RNA modification machinery reveal tissue- and cancer-specific signatures. Genome Biol. 2020;21: 97. doi:10.1186/s13059-020-02009-z

7. Saliminejad K, Khorram Khorshid HR, Soleymani Fard S, Ghaffari SH. An overview of microRNAs: Biology, functions, therapeutics, and analysis methods. J Cell Physiol. 2019;234: 5451–5465. doi:10.1002/jcp.27486

8. Kao H-W, Pan C-Y, Lai C-H, Wu C-W, Fang W-L, Huang K-H, et al. Urine miR-21-5p as a potential non-invasive biomarker for gastric cancer. Oncotarget. 2017;8: 56389–56397. doi:10.18632/oncotarget.16916

9. Shi R, Wang P-Y, Li X-Y, Chen J-X, Li Y, Zhang X-Z, et al. Exosomal levels of miRNA-21 from cerebrospinal fluids associated with poor prognosis and tumor recurrence of glioma patients. Oncotarget. 2015;6: 26971–26981. doi:10.18632/oncotarget.4699

10. Peng Q, Zhang X, Min M, Zou L, Shen P, Zhu Y. The clinical role of microRNA-21 as a promising biomarker in the diagnosis and prognosis of colorectal cancer: a systematic review and meta-analysis. Oncotarget. 2017;8: 44893–44909. doi:10.18632/oncotarget.16488

11. Khalid U, Ablorsu E, Szabo L, Jenkins RH, Bowen T, Chavez R, et al. MicroRNA-21 (miR-21) expression in hypothermic machine perfusate may be predictive of early outcomes in kidney transplantation. Clin Transplant. 2016;30: 99–104. doi:10.1111/ctr.12679

12. Elbehidy RM, Youssef DM, El-Shal AS, Shalaby SM, Sherbiny HS, Sherief LM, et al. MicroRNA-21 as a novel biomarker in diagnosis and response to therapy in asthmatic children. Mol Immunol. 2016;71: 107–114. doi:10.1016/j.molimm.2015.12.015

13. Olivieri F, Spazzafumo L, Bonafè M, Recchioni R, Prattichizzo F, Marcheselli F, et al. MiR-21-5p and miR-126a-3p levels in plasma and circulating angiogenic cells: relationship with type 2 diabetes complications. Oncotarget. 2015;6: 35372–35382. doi:10.18632/oncotarget.6164

14. Migita K, Komori A, Kozuru H, Jiuchi Y, Nakamura M, Yasunami M, et al. Circulating microRNA Profiles in Patients with Type-1 Autoimmune Hepatitis. PLoS One. 2015;10: e0136908. doi:10.1371/journal.pone.0136908

15. Zendjabil M, Favard S, Tse C, Abbou O, Hainque B. [The microRNAs as biomarkers: What prospects?]. C R Biol. 2017;340: 114–131. doi:10.1016/j.crvi.2016.12.001

16. Armand-Labit V, Pradines A. Circulating cell-free microRNAs as clinical cancer biomarkers. Biomol Concepts. 2017;8: 61–81. doi:10.1515/bmc-2017-0002

17. He J-H, Li Y-G, Han Z-P, Zhou J-B, Chen W-M, Lv Y-B, et al. The CircRNA-ACAP2/Hsa-miR-21-5p/ Tiam1 Regulatory Feedback Circuit Affects the Proliferation, Migration, and Invasion of Colon Cancer SW480 Cells. Cell Physiol Biochem. 2018;49: 1539–1550. doi:10.1159/000493457

18. Tse J, Pierce T, Carli ALE, Alorro MG, Thiem S, Marcusson EG, et al. Onco-miR-21 Promotes Stat3-Dependent Gastric Cancer Progression. Cancers (Basel). 2022;14: 264. doi:10.3390/cancers14020264

19. Jiang J, Wang X, Lu J. PWRN1 Suppressed Cancer Cell Proliferation and Migration in Glioblastoma by Inversely Regulating hsa-miR-21-5p. Cancer Manag Res. 2020;12: 5313–5322. doi:10.2147/CMAR.S250166

20. Yu W, Zhu K, Wang Y, Yu H, Guo J. Overexpression of miR-21-5p promotes proliferation and invasion of colon adenocarcinoma cells through targeting CHL1. Molecular Medicine. 2018;24: 36. doi:10.1186/s10020-018-0034-5

21. Lim SL, Qu ZP, Kortschak RD, Lawrence DM, Geoghegan J, Hempfling A-L, et al. HENMT1 and piRNA Stability Are Required for Adult Male Germ Cell Transposon Repression and to Define the Spermatogenic Program in the Mouse. PLoS Genet. 2015;11: e1005620. doi:10.1371/journal.pgen.1005620

22. Hempfling AL, Lim SL, Adelson DL, Evans J, O’Connor AE, Qu ZP, et al. Expression patterns of HENMT1 and PIWIL1 in human testis: implications for transposon expression. Reproduction. 2017;154: 363–374. doi:10.1530/REP-16-0586

23. Liang H, Jiao Z, Rong W, Qu S, Liao Z, Sun X, et al. 3’-Terminal 2’-O-methylation of lung cancer miR-21-5p enhances its stability and association with Argonaute 2. Nucleic Acids Res. 2020;48: 7027–7040. doi:10.1093/nar/gkaa504

24. Piuco R, Galante PAF. piRNAdb: A piwi-interacting RNA database. bioRxiv; 2021. p. 2021.09.21.461238. doi:10.1101/2021.09.21.461238

25. Gainetdinov IV, Skvortsova YV, Kondratieva SA, Klimov A, Tryakin AA, Azhikina TL. Assessment of piRNA biogenesis and function in testicular germ cell tumors and their precursor germ cell neoplasia in situ. BMC Cancer. 2018;18: 20. doi:10.1186/s12885-017-3945-6

26. Roovers EF, Rosenkranz D, Mahdipour M, Han C-T, He N, Chuva de Sousa Lopes SM, et al. Piwi proteins and piRNAs in mammalian oocytes and early embryos. Cell Rep. 2015;10: 2069–2082. doi:10.1016/j.celrep.2015.02.062

27. Williams Z, Morozov P, Mihailovic A, Lin C, Puvvula PK, Juranek S, et al. Discovery and Characterization of piRNAs in the Human Fetal Ovary. Cell Rep. 2015;13: 854–863. doi:10.1016/j.celrep.2015.09.030

28. Sai Lakshmi S, Agrawal S. piRNABank: a web resource on classified and clustered Piwi-interacting RNAs. Nucleic Acids Res. 2008;36: D173–D177. doi:10.1093/nar/gkm696

29. Das PP, Bagijn MP, Goldstein LD, Woolford JR, Lehrbach NJ, Sapetschnig A, et al. Piwi and piRNAs Act Upstream of an Endogenous siRNA Pathway to suppress Tc3 Transposon Mobility in the Caenorhabditis elegans germline. Mol Cell. 2008;31: 79–90. doi:10.1016/j.molcel.2008.06.003

30. Grimson A, Srivastava M, Fahey B, Woodcroft BJ, Chiang HR, King N, et al. Early origins and evolution of microRNAs and Piwi-interacting RNAs in animals. Nature. 2008;455: 1193–1197. doi:10.1038/nature07415

31. Huang Y, Ji L, Huang Q, Vassylyev DG, Chen X, Ma J-B. Structural insights into mechanisms of the small RNA methyltransferase HEN1. Nature. 2009;461: 823–827. doi:10.1038/nature08433

32. Yu B, Yang Z, Li J, Minakhina S, Yang M, Padgett RW, et al. Methylation as a Crucial Step in Plant microRNA Biogenesis. Science. 2005;307: 932–935. doi:10.1126/science.1107130

33. Yang Z, Ebright YW, Yu B, Chen X. HEN1 recognizes 21-24 nt small RNA duplexes and deposits a methyl group onto the 2’ OH of the 3’ terminal nucleotide. Nucleic Acids Res. 2006;34: 667–675. doi:10.1093/nar/gkj474

34. Horwich MD, Li C, Matranga C, Vagin V, Farley G, Wang P, et al. The Drosophila RNA methyltransferase, DmHen1, modifies germline piRNAs and single-stranded siRNAs in RISC. Curr Biol. 2007;17: 1265–1272. doi:10.1016/j.cub.2007.06.030

35. Saito K, Sakaguchi Y, Suzuki T, Suzuki T, Siomi H, Siomi MC. Pimet, the Drosophila homolog of HEN1, mediates 2′-O-methylation of Piwi-interacting RNAs at their 3′ ends. Genes Dev. 2007;21: 1603–1608. doi:10.1101/gad.1563607

36. Chan CM, Zhou C, Brunzelle JS, Huang RH. Structural and biochemical insights into 2′-O-methylation at the 3′-terminal nucleotide of RNA by Hen1. PNAS. 2009;106: 17699–17704. doi:10.1073/pnas.0907540106

37. Peng L, Zhang F, Shang R, Wang X, Chen J, Chou JJ, et al. Identification of substrates of the small RNA methyltransferase Hen1 in mouse spermatogonial stem cells and analysis of its methyl-transfer domain. J Biol Chem. 2018;293: 9981–9994. doi:10.1074/jbc.RA117.000837

38. Kirino Y, Mourelatos Z. The mouse homolog of HEN1 is a potential methylase for Piwi-interacting RNAs. RNA. 2007;13: 1397–1401. doi:10.1261/rna.659307

39. Kamminga LM, Luteijn MJ, den Broeder MJ, Redl S, Kaaij LJT, Roovers EF, et al. Hen1 is required for oocyte development and piRNA stability in zebrafish. EMBO J. 2010;29: 3688–3700. doi:10.1038/emboj.2010.233

40. Huang RH. Unique 2′-O-Methylation by Hen1 in Eukaryotic RNA Interference and Bacterial RNA Repair. Biochemistry. 2012;51: 4087–4095. doi:10.1021/bi300497x

41. Zhao LN, Kaldis P. Cascading proton transfers are a hallmark of the catalytic mechanism of SAM-dependent methyltransferases. FEBS Letters. 2020;594: 2128–2139. doi:10.1002/1873-3468.13799

42. Fiser A, Do RK, Sali A. Modeling of loops in protein structures. Protein Sci. 2000;9: 1753–1773. doi:10.1110/ps.9.9.1753

43. Singh N, Warshel A. Absolute binding free energy calculations: On the accuracy of computational scoring of protein–ligand interactions. Proteins: Structure, Function, and Bioinformatics. 2010;78: 1705–1723. doi:10.1002/prot.22687

44. Sham YY, Chu ZT, Warshel A. Consistent Calculations of pKa’s of Ionizable Residues in Proteins: Semi-microscopic and Microscopic Approaches. J Phys Chem B. 1997;101: 4458–4472. doi:10.1021/jp963412w

45. King G, Warshel A. A surface constrained all-atom solvent model for effective simulations of polar solutions. J Chem Phys. 1989;91: 3647–3661. doi:10.1063/1.456845

46. Yoon H, Zhao LN, Warshel A. Exploring the Catalytic Mechanism of Cas9 Using Information Inferred from Endonuclease VII. ACS Catal. 2019;9: 1329–1336. doi:10.1021/acscatal.8b04324

47. Zhao LN, Mondal D, Warshel A. Exploring alternative catalytic mechanisms of the Cas9 HNH domain. Proteins: Structure, Function, and Bioinformatics. 2020;88: 260–264. doi:10.1002/prot.25796

48. Lee FS, Chu ZT, Warshel A. Microscopic and semimicroscopic calculations of electrostatic energies in proteins by the POLARIS and ENZYMIX programs. Journal of Computational Chemistry. 1993;14: 161–185. doi:10.1002/jcc.540140205

49. Frisch MJ, Trucks GW, Schlegel HB, Scuseria GE, Robb MA, Cheeseman JR, et al. Gaussian^∼^16 Revision C.01. 2016.

50. Thaplyal P, Bevilacqua PC. Experimental Approaches for Measuring pKa’s in RNA and DNA. Methods Enzymol. 2014;549: 189–219. doi:10.1016/B978-0-12-801122-5.00009-X

51. Li H, Robertson AD, Jensen JH. Very fast empirical prediction and rationalization of protein pKa values. Proteins. 2005;61: 704–721. doi:10.1002/prot.20660

52. Mordasini T, Curioni A, Andreoni W. Why Do Divalent Metal Ions Either Promote or Inhibit Enzymatic Reactions? Journal of Biological Chemistry. 2003;278: 4381–4384. doi:10.1074/jbc.C200664200

53. Ayadi L, Galvanin A, Pichot F, Marchand V, Motorin Y. RNA ribose methylation (2′-O-methylation): Occurrence, biosynthesis and biological functions. Biochimica et Biophysica Acta (BBA) - Gene Regulatory Mechanisms. 2019;1862: 253–269. doi:10.1016/j.bbagrm.2018.11.009

54. Bock CW, Katz AK, Markham GD, Glusker JP. Manganese as a Replacement for Magnesium and Zinc: Functional Comparison of the Divalent Ions. J Am Chem Soc. 1999;121: 7360–7372. doi:10.1021/ja9906960

55. Kretsinger RH. Magnesium in Biological Systems. In: Kretsinger RH, Uversky VN, Permyakov EA,editors. Encyclopedia of Metalloproteins. New York, NY: Springer; 2013. pp. 1250–1255. doi:10.1007/978-1-4614-1533-6_258

56. Tang S, Yang JJ. Magnesium Binding Sites in Proteins. In: Kretsinger RH, Uversky VN, Permyakov EA, editors. Encyclopedia of Metalloproteins. New York, NY: Springer; 2013. pp. 1243–1250. doi:10.1007/978-1-4614-1533-6_257

57. Zheng H, Cooper DR, Porebski PJ, Shabalin IG, Handing KB, Minor W. CheckMyMetal: a macromolecular metal-binding validation tool. Acta Crystallogr D Struct Biol. 2017;73: 223–233. doi:10.1107/S2059798317001061

58. Zheng H, Chordia MD, Cooper DR, Chruszcz M, Müller P, Sheldrick GM, et al. Validation of metal-binding sites in macromolecular structures with the CheckMyMetal web server. Nat Protoc. 2014;9: 156–170. doi:10.1038/nprot.2013.172

59. Jain R, Shuman S. Bacterial Hen1 is a 3′ terminal RNA ribose 2′-O-methyltransferase component of a bacterial RNA repair cassette. RNA. 2010;16: 316–323. doi:10.1261/rna.1926510

60. Maguire ME, Cowan JA. Magnesium chemistry and biochemistry. Biometals. 2002;15: 203–210. doi:10.1023/A:1016058229972

61. Diebler H, Eigen M, Ilgenfritz G, Maass G, Winkler R. Kinetics and mechanism of reactions of main group metal ions with biological carriers. Pure and Applied Chemistry. 1969;20: 93–116. doi:10.1351/pac196920010093

62. Vilkaitis G, Plotnikova A, Klimasauskas S. Kinetic and functional analysis of the small RNA methyltransferase HEN1: The catalytic domain is essential for preferential modification of duplex RNA. RNA. 2010;16: 1935–1942. doi:10.1261/rna.2281410

63. Jain R, Shuman S. Active site mapping and substrate specificity of bacterial Hen1, a manganese-dependent 3′ terminal RNA ribose 2′O-methyltransferase. RNA. 2011;17: 429–438. doi:10.1261/rna.2500711

64. Yao J, Xie M, Ma X, Song J, Wang Y, Xue X. PIWI-interacting RNAs in cancer: Biogenesis, function, and clinical significance. Frontiers in Oncology. 2022;12. Available: https://www.frontiersin.org/articles/10.3389/fonc.2022.965684

